# Matrix Matters: Context-Driven Metabolic Shifts in *Bacillus cereus* and *Bacillus subtilis*

**DOI:** 10.1101/2025.10.06.680604

**Authors:** Emilie Viktoria Skriver, Tessa Suzanne Canoy, Yuandong Sha, Morten Arendt Rasmussen, Bekzod Khakimov, Susanne Knøchel, Henriette Lyng Røder

**Affiliations:** Department of Food Science, University of Copenhagen, Copenhagen, Denmark; Copenhagen Studies on Asthma in Childhood, Herlev-Gentofte Hospital, University of Copenhagen, Copenhagen, Denmark

**Keywords:** *Bacillus cereus*, *Bacillus subtilis*, isothermal calorimetry, microbial metabolism, cereulide, food matrix, plant-based food, metabolomics

## Abstract

Spore-forming *Bacillus* species, including pathogenic *Bacillus cereus* and spoilage-associated *Bacillus subtilis*, are major contributors to foodborne illness and product degradation. Understanding their metabolic behaviour in diverse food matrices is essential for improving risk assessment, spoilage prediction, and fermentation control. This study integrates isothermal microcalorimetry and targeted metabolomics to characterize the metabolic activity of *B. cereus* and *B. subtilis* in five nutrient sources: Brain Heart Infusion (BHI) medium, oat drink, milk, pea hydrolysate, and a combined oat-pea matrix. Metabolic heat production was monitored for 24 hours at 30°C. In BHI, *B. cereus* exhibited a shorter lag phase (mean ± sd: 4.3 hours ± 0.8) than *B. subtilis* (7.9 hours ± 1.0) but produced less total heat. Across all food matrices, *B. subtilis* consistently generated more heat. The oat-pea matrix supported the highest calorimetric growth rates, surpassing oat or pea alone, and showed sugar depletion and accumulation of organic acids, indicating enhanced carbohydrate metabolism. Free amino acid release was matrix- and species-specific: *B. subtilis* had increased levels in oat, while *B. cereus* did so in pea. While *B. cereus* was metabolically active in all matrices, cereulide levels were matrix-dependent: 47.3 ± 1.7 ng/mL in oat, 3.0 ± 0.1 ng/mL in oat–pea, and undetectable in pea. These findings reveal clade-specific and matrix-driven metabolic strategies. This is the first study to combine calorimetry and metabolomics to evaluate *Bacillus* activity in plant-based and dairy matrices. This approach enhances our understanding of microbial physiology in complex food systems and provides a foundation for developing targeted strategies to improve food safety, stability, and product design.

## 1. Introduction

Microbial contamination is a key concern in the food industry (Snyder et al., 2024). In 2023, the United Nations estimated that 19% of all food available at the consumer level was wasted, which has a significant impact on the economy, environment, and public health (UNEP, 2024). The Gram-positive, spore-forming, facultative aerobe *Bacillus* spp. are ubiquitous in nature and include species such as *B. cereus* and *B. subtilis*. *Bacillus* species frequently enter food production system through raw ingredients (Ehling-Schulz et al., 2015). Their highly resistant spores complicate control with conventional heat, acid, and cleaning practices (Earl et al., 2008; Rahnama et al., 2023).

*Bacillus cereus* is an important foodborne pathogen, capable of causing diarrhoea due to enterotoxins or inducing vomiting due to the pre-formed heat- and acid-stable emetic toxin, cereulide (Ehling-Schulz et al., 2004). The *B. cereus* group, also known as *B. cereus* sensu lato, contains many genetically closely related species, including *B. cereus, B. mycoides,* and *B. thuringiensis*. *B. cereus* taxonomy has undergone many changes over the years, especially after the introduction of genome sequencing (Carroll et al., 2022).

Foodborne disease caused by *B. cereus* is believed to be grossly underreported. Nevertheless, in 2024, *B. cereus* toxins (both emetic and enterotoxins) caused 127 registered foodborne outbreaks, 3315 cases of illness, 49 hospitalizations, and 9 deaths in the EU and UK alone (EFSA, 2025). *B. cereus* is typically isolated from plant materials (Messelhäusser et al., 2014), and the expanding use of plant-based ingredients in food production increases the relevance of monitoring this pathogen in plant matrices (Kyrylenko et al., 2023). Cereulide warrants particular attention, as its exceptional heat and acid stability allow it to persist through processing and cause intoxication even at very low concentrations (Rajkovic et al., 2008).

*B. subtilis* is a key driver of many traditional fermentations based on legumes, seeds, and tubers (Li et al., 2023). A high occurrence of *B. cereus* has been observed in *B. subtilis-*fermented food products, posing a significant concern for food safety (Akanni et al., 2018; Owusu-Kwarteng et al., 2022). In some cases, *B. subtilis* is also associated with food spoilage, such as slime formation in rice products and ropiness in bread (Dong et al., 2023; Valerio et al., 2008). Despite frequent co-occurrence, species-resolved metabolic characteristics and growth dynamics of *B. cereus* and *B. subtilis* in food-relevant matrices remain insufficiently characterised. Elucidating these species-specific responses is essential for predicting their roles in fermentation, spoilage, and food safety risks in plant-based foods.

An attractive tool for fast detection of microbial activity is isothermal microcalorimetry (IMC). IMC is a non-invasive, real-time readout of heat flow generated by the metabolic activity of biological systems, and is particularly useful in opaque or complex matrices where optical methods are not applicable. IMC can serve as a quantitative proxy for active biomass because heat flow scales with metabolically active cell numbers, with strong correlations reported between calorimetric output and optical or culture-based biomass measurements (Braissant et al., 2015a; Braissant et al., 2010). Importantly, heat flow captures total metabolic activity and therefore reports on active biomass without distinguishing whether changes come from cell numbers or per-cell metabolic rates. IMC equipment has become less costly and has demonstrated improved sensitivity and accuracy (Braissant et al., 2010). IMC has recently been successfully applied to improve bacterial detection time in a clinical context (Cichos et al., 2023) and to investigate the metabolic activity of bacteria living in biofilms (Lichtenberg et al., 2022). Despite this potential for non-invasive microbial testing of foods, IMC has not been applied to monitor the growth of pathogenic and non-pathogenic *Bacillus* spp. in food relevant matrices.

While IMC provides an integrated measure of microbial metabolic activity, it offers limited specificity regarding the biochemical processes underlying bacterial metabolism. Monitoring changes in key metabolites such as sugars, organic acids, and amino acids enables a deeper understanding of microbial nutrient assimilation while also revealing biochemical changes in the food matrices that are relevant to safety and quality. Sugars serve as primary energy sources for microbial growth, while organic acids are key intermediates in carbon metabolism (Schilling et al., 2007). Free amino acids are an important nitrogen source and can be released either through proteolytic activity or produced through *de novo* synthesis (Sonenshein, 2007). Furthermore, quantifying the heat-stable emetic toxin cereulide is essential for evaluating the safety risks associated with food products contaminated by *B. cereus*.

To address these needs, we integrate IMC with targeted metabolomics to provide a comprehensive and mechanistic understanding of *Bacillus* physiology in food-relevant matrices. Previous IMC applications in food science have largely focused on microbial growth dynamics, such as lactic acid bacteria performance in sourdough (Mihhalevski et al., 2011), the effect of glucose depletion on growth (Kabanova et al., 2013), shelf life of carrot juice (Alklint et al., 2005), and yeast inhibition by alcohol (Antoce et al., 1997). Here, we specifically investigate the metabolic heat production of *B. cereus* and *B. subtilis* in a rich laboratory medium and in dairy- and plant-based liquid matrices (hereafter referred to as matrices), while simultaneously quantifying sugars, organic acids, free amino acids, and cereulide at selected time points (Figure 1). This dual approach of combining IMC with metabolomics enables a novel perspective on *Bacillus* activity in food systems, providing insights that go beyond growth kinetics to inform both food safety and quality management.

**Figure 1:**
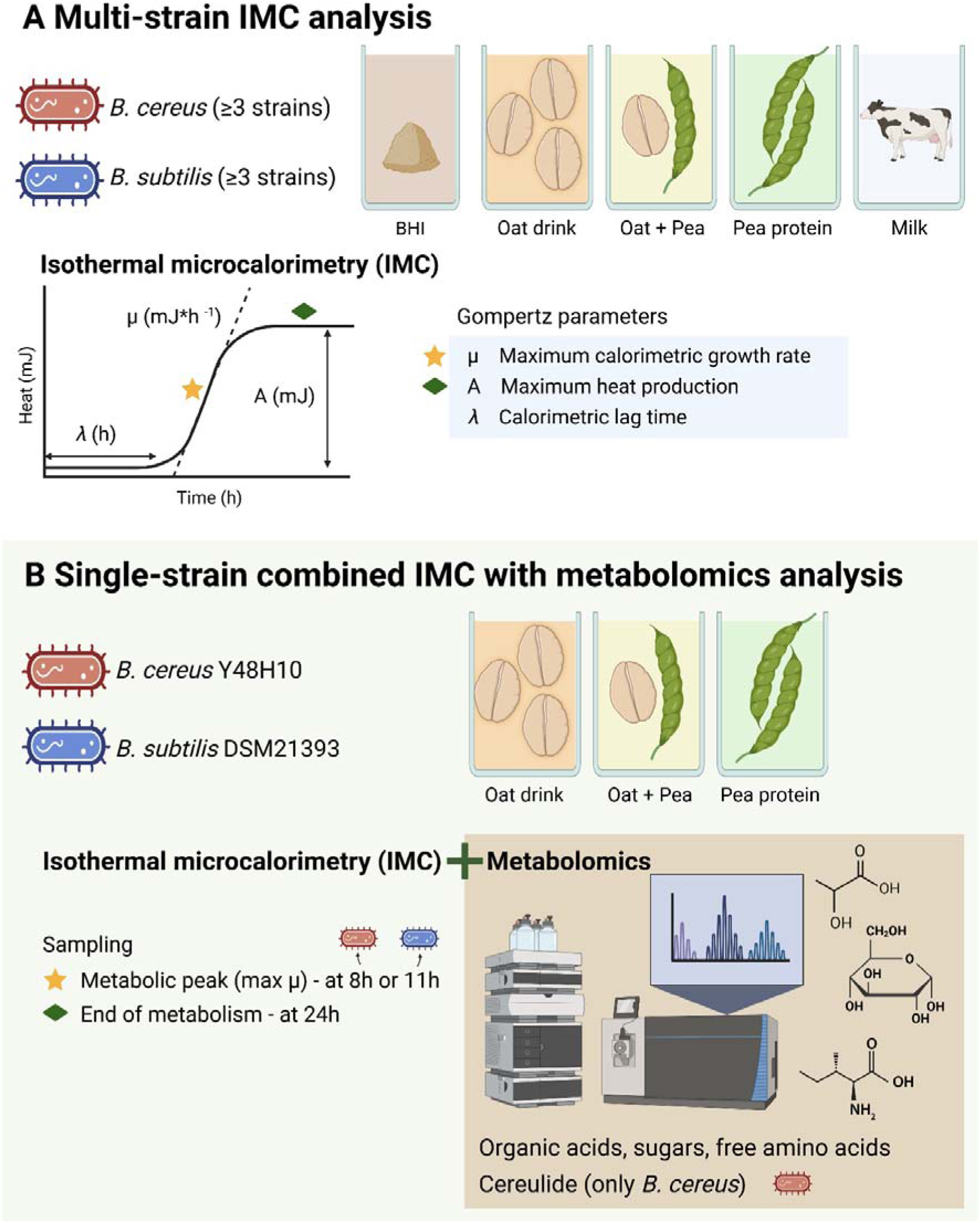
Experimental workflow combining isothermal microcalorimetry (IMC) and targeted metabolomics. IMC was applied to monitor metabolic activity of *B. cereus* and *B. subtilis* in different food-relevant matrices, and to define sampling points for metabolite analysis. (A) Multi-strain IMC characterization: Sixteen *B. cereus* and eight *B. subtilis* strains were cultivated in BHI broth to characterize heat flow dynamics over 20 h. A subset of three strains per species was further assessed in four liquid food matrices (milk, oat, pea, oat–pea blend) for 24 h. (B) Integrated IMC– metabolomics analysis: One representative strain from each species (*B. cereus* Y48H10 and *B. subtilis* DSM21393) was cultivated in oat, pea and oat-pea matrices. IMC guided sampling at two time points: peak metabolic activity at maximum calorimetric growth rate (star) and end-of-metabolism at maximum heat production (diamond). At these points, pooled samples were analysed for sugars, organic acids, and free amino acids; cereulide was quantified in *B. cereus* only. Figure created in BioRender https://BioRender.com/obt15z8.

## 2. Materials and methods

### 2.1 Bacterial strains

Sixteen strains from the *B. cereus* group and eight *B. subtilis* strains were selected based on diversity in toxicity, genetic characteristics, and origin (Table 1). Five of the included *B. subtilis* strains were previously isolated from traditional fermentations in Ghana and Burkina Faso (Kpikpi et al., 2014; Parkouda et al., 2010; Wiedenbein et al., 2023). Two of the *B. cereus* strains were isolated from African fermented condiments (Thorsen et al., 2010). Strains were maintained in a -80°C glycerol stock and streaked onto Brain Heart Infusion (BHI) agar plates before being cultured in BHI broth.

**TABLE 1:**
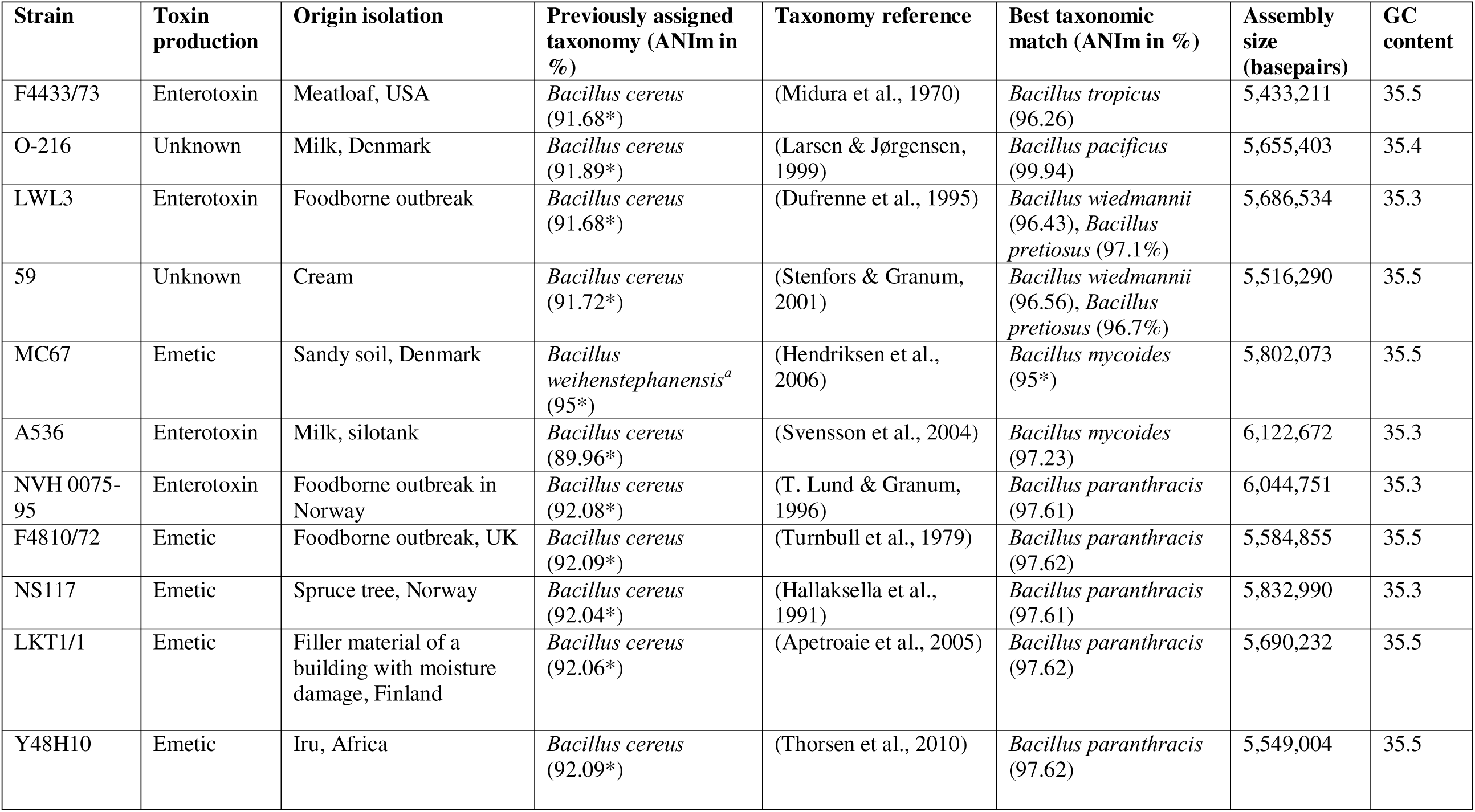
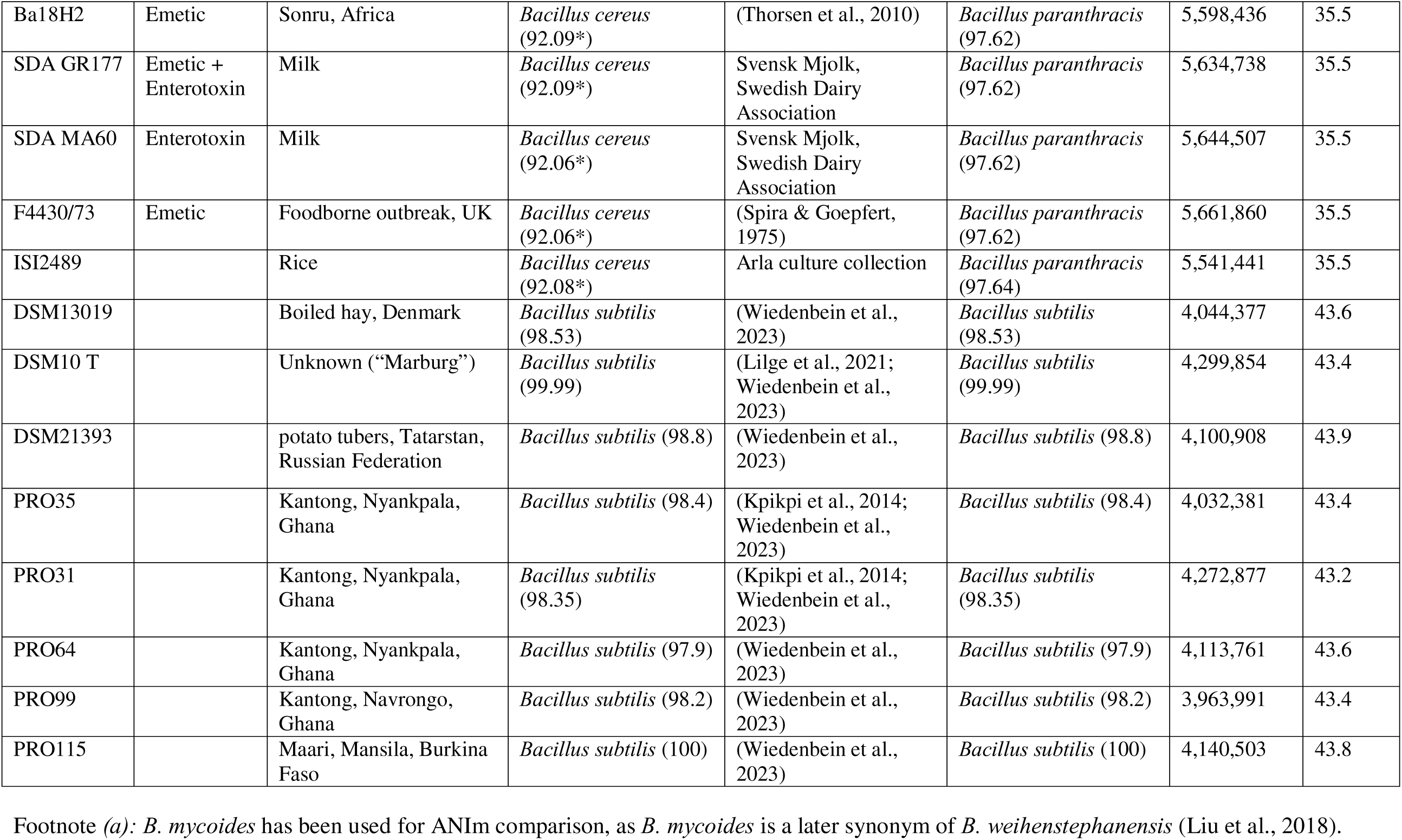
Characteristics of the *Bacillus* strains used in this study, based on sequencing and ANIm data for the *Bacillus cereus* group obtained by Skriver et al. (2026) and for the *Bacillus subtilis* strains obtained by Wiedenbein et al. (2023). Asterisk (*) denotes values below species threshold (<96% ANIm). The letter T refers to the type strain. Toxin production for *B. cereus* isolates refers to unknown toxin production, literature-reported production of exclusively cereulide,, enterotoxin or both cereulide and enterotoxin.

| Strain | Toxin production | Origin isolation | Previously assigned taxonomy (ANIm in %) | Taxonomy reference | Best taxonomic match (ANIm in %) | Assembly size (basepairs) | GC content |
| --- | --- | --- | --- | --- | --- | --- | --- |
| F4433/73 | Enterotoxin | Meatloaf, USA | *Bacillus cereus* (91.68*) | (Midura et al., 1970) | *Bacillus tropicus* (96.26) | 5,433,211 | 35.5 |
| O-216 | Unknown | Milk, Denmark | *Bacillus cereus* (91.89*) | (Larsen & Jørgensen, 1999) | *Bacillus pacificus* (99.94) | 5,655,403 | 35.4 |
| LWL3 | Enterotoxin | Foodborne outbreak | *Bacillus cereus* (91.68*) | (Dufrenne et al., 1995) | *Bacillus wiedmannii* (96.43), *Bacillus pretiosus* (97.1%) | 5,686,534 | 35.3 |
| 59 | Unknown | Cream | *Bacillus cereus* (91.72*) | (Stenfors & Granum, 2001) | *Bacillus wiedmannii* (96.56), *Bacillus pretiosus* (96.7%) | 5,516,290 | 35.5 |
| MC67 | Emetic | Sandy soil, Denmark | *Bacillus weihenstephanensis*<sup>a</sup> (95*) | (Hendriksen et al., 2006) | *Bacillus mycoides* (95*) | 5,802,073 | 35.5 |
| A536 | Enterotoxin | Milk, silotank | *Bacillus cereus* (89.96*) | (Svensson et al., 2004) | *Bacillus mycoides* (97.23) | 6,122,672 | 35.3 |
| NVH 0075-95 | Enterotoxin | Foodborne outbreak in Norway | *Bacillus cereus* (92.08*) | (T. Lund & Granum, 1996) | *Bacillus paranthracis* (97.61) | 6,044,751 | 35.3 |
| F4810/72 | Emetic | Foodborne outbreak, UK | *Bacillus cereus* (92.09*) | (Turnbull et al., 1979) | *Bacillus paranthracis* (97.62) | 5,584,855 | 35.5 |
| NS117 | Emetic | Spruce tree, Norway | *Bacillus cereus* (92.04*) | (Hallaksella et al., 1991) | *Bacillus paranthracis* (97.61) | 5,832,990 | 35.3 |
| LKT1/1 | Emetic | Filler material of a building with moisture damage, Finland | *Bacillus cereus* (92.06*) | (Apetroaie et al., 2005) | *Bacillus paranthracis* (97.62) | 5,690,232 | 35.5 |
| Y48H10 | Emetic | Iru, Africa | *Bacillus cereus* (92.09*) | (Thorsen et al., 2010) | *Bacillus paranthracis* (97.62) | 5,549,004 | 35.5 |
| Ba18H2 | Emetic | Sonru, Africa | *Bacillus cereus* (92.09*) | (Thorsen et al., 2010) | *Bacillus paranthracis* (97.62) | 5,598,436 | 35.5 |
| SDA GR177 | Emetic + Enterotoxin | Milk | *Bacillus cereus* (92.09*) | Svensk Mjolk, Swedish Dairy Association | *Bacillus paranthracis* (97.62) | 5,634,738 | 35.5 |
| SDA MA60 | Enterotoxin | Milk | *Bacillus cereus* (92.06*) | Svensk Mjolk, Swedish Dairy Association | *Bacillus paranthracis* (97.62) | 5,644,507 | 35.5 |
| F4430/73 | Emetic | Foodborne outbreak, UK | *Bacillus cereus* (92.06*) | (Spira & Goepfert, 1975) | *Bacillus paranthracis* (97.62) | 5,661,860 | 35.5 |
| ISI2489 |  | Rice | *Bacillus cereus* (92.08*) | Arla culture collection | *Bacillus paranthracis* (97.64) | 5,541,441 | 35.5 |
| DSM13019 |  | Boiled hay, Denmark | *Bacillus subtilis* (98.53) | (Wiedenbein et al., 2023) | *Bacillus subtilis* (98.53) | 4,044,377 | 43.6 |
| DSM10 T |  | Unknown (“Marburg”) | *Bacillus subtilis* (99.99) | (Lilge et al., 2021; Wiedenbein et al., 2023) | *Bacillus subtilis* (99.99) | 4,299,854 | 43.4 |
| DSM21393 |  | potato tubers, Tatarstan, Russian Federation | *Bacillus subtilis* (98.8) | (Wiedenbein et al., 2023) | *Bacillus subtilis* (98.8) | 4,100,908 | 43.9 |
| PRO35 |  | Kantong, Nyankpala, Ghana | *Bacillus subtilis* (98.4) | (Kpikpi et al., 2014; Wiedenbein et al., 2023) | *Bacillus subtilis* (98.4) | 4,032,381 | 43.4 |
| PRO31 |  | Kantong, Nyankpala, Ghana | *Bacillus subtilis* (98.35) | (Kpikpi et al., 2014; Wiedenbein et al., 2023) | *Bacillus subtilis* (98.35) | 4,272,877 | 43.2 |
| PRO64 |  | Kantong, Nyankpala, Ghana | *Bacillus subtilis* (97.9) | (Wiedenbein et al., 2023) | *Bacillus subtilis* (97.9) | 4,113,761 | 43.6 |
| PRO99 |  | Kantong, Navrongo, Ghana | *Bacillus subtilis* (98.2) | (Wiedenbein et al., 2023) | *Bacillus subtilis* (98.2) | 3,963,991 | 43.4 |
| PRO115 |  | Maari, Mansila, Burkina Faso | *Bacillus subtilis* (100) | (Wiedenbein et al., 2023) | *Bacillus subtilis* (100) | 4,140,503 | 43.8 |

Footnote (a): *B. mycoides* has been used for ANIm comparison, as *B. mycoides* is a later synonym of *B. weihenstephanensis* (Liu et al., 2018).

### 2.2 Whole genome sequencing

*B. subtilis* whole genomes included in this work were previously characterized in Wiedenbein et al. (2023), while the *B. cereus* whole genomes were obtained from Skriver et al. (2026).

### 2.3 Phylogenetic tree construction

Phylogenetic trees were constructed with the up-to-date bacterial core gene (UBCG) method based on the concatenation of 92 core genes found in all taxonomic ranks of Bacteria (Na et al., 2018). FastTree was selected to generate the maximum-likelihood phylogenetic trees (Price et al., 2010). The phylogenetic trees were visualized with Molecular Evolutionary Genetics Analysis version 11 software (MEGA 11) (Tamura et al., 2021). *Halobacillus halophilus* DSM 2266 (type strain) (GenBank: GCA_000284515.1) was selected as the outgroup to root the trees.

### 2.4 Isothermal calorimetric assessment of *B. cereus* and *B. subtilis* in BHI medium

All sixteen *B. cereus* and eight *B. subtilis* strains (Table 1) were cultivated in BHI broth (Merck). To measure the heat flow of the *Bacillus* strains in BHI medium, the protocol of Beilharz et al. (2023) was followed with a few modifications. Strains were cultured in BHI broth overnight (30°C, 200 rpm) and then diluted with BHI broth to an optical density at 600 nm (OD_600_) of 0.2. The inoculum was mixed with fresh BHI medium at a 1% inoculation rate to a final volume of 200 µL, corresponding to an initial concentration of 10^6^ CFU/mL. The inoculated mixtures were transferred to plastic inserts (calWells™, Symcel AB, Sweden) in the titanium vials of a 48-well calPlate™ (Symcel AB, Sweden). The wells were sealed using a torque wrench set on 40 cNm force to ensure a closed system for monitoring heat exchange. Non-inoculated BHI medium was dispensed into 16 reference vials, one for every two of the 32 sample vials, to account for BHI-dependent effects on system-generated heat. The vials were pre-incubated for 60 min to reach temperature equilibrium. The plates were then incubated in the Calscreener (Symcel AB, Sweden) at 30°C for 20h.

### 2.5 Isothermal calorimetric assessment of *B. cereus* and *B. subtilis* in food matrices

Ultra-high temperature treated (UHT) whole milk (Geia Food A/S, Czech Republic) and UHT oat drink (Naturli, Denmark) were purchased commercially in Danish supermarkets. For the mixed oat-pea matrix, 5% w/w pea protein hydrolysate (NUTRALYS® PEA F853M-EXP, Roquette, France) was transferred to the store-bought oat drink and autoclaved (121°C, 20 min). For the pea matrix, 5% w/w was added to MilliQ water and autoclaved (121°C, 20 min). The pea and oat-pea matrices were autoclaved prior to inoculation to solubilize the pea hydrolysate and eliminate contaminants present in the pea isolate. In contrast, the UHT oat drink was not further autoclaved, as autoclaving caused growth inhibition by *B. cereus* in preliminary experiments. These differences in thermal processing were considered when interpreting metabolic outcomes.

A selection of three *B. cereus* strains (Y48H10, SDA GR177, MC67) and three *B. subtilis* strains (PRO99, DSM10, DSM21393) were cultivated in the food matrices. Cells were cultured in BHI broth overnight (30°C, 200 rpm), centrifuged at 4500 *g* for 12 minutes, and washed twice in 0.9% NaCl solution. Washed cells were adjusted to OD_600_ of 0.2 before being transferred with an inoculation rate of 1% and a total volume of 200 µL into the different food matrices. Similarly to section 2.4, the inoculated mixtures were transferred to plastic inserts in the titanium vials of the 48-well calPlate and pre-incubated for 60 min to reach temperature equilibrium. Non-inoculated liquid food matrices were dispensed into 16 reference vials, one for every two of the 32 sample vials, to account for matrix-dependent effects on system-generated heat. The calPlate was then incubated in the Calscreener at 30°C for 24 hours.

### 2.6 Cultivation of selected strains in oat, pea, and oat-pea matrices for metabolic profiling

One strain from each species (*B. cereus* Y48H10 and *B. subtilis* DSM21393) was selected for in-depth metabolic profiling. Washed cells were prepared as in section 2.5 and then inoculated in oat, pea, and oat-pea matrices. The initial pH of each food matrix was measured using a Radiometer PHM250 pH meter (Radiometer Analytical, Denmark). Two biological replicates, each consisting of four technical replicates, were used. Biological replicates were defined as individual colonies streaked from the same BHI plate, while technical replicates originated from the same biological replicate after two rounds of pre-cultivation on BHI broth. Samples were taken at maximum calorimetric growth rate, approximately 8 hours for *B. cereus* and 11 hours for *B. subtilis*, and at the end-metabolic point, 24 hours for both *B. cereus* and *B. subtilis*. Directly after sampling, technical replicates were pooled to obtain a final volume of approximately 800 μL. This pooled sample was analysed for sugars and organic acids (sections 2.7), free amino acids (section 2.8), and the *B. cereus* samples were screened for cereulide (section 2.9).

### 2.7 Quantification of simple sugars and organic acids

Pooled technical replicates of food samples (section 2.6) were centrifuged at 12000 *g* for 15 minutes at 4°C, and the supernatant was separated and filtered through PTFE membranes with a pore size of 0.45 µm directly into the HPLC vials and stored at -21°C until analysis. An Agilent 1260 Infinity II LC system with refractive index detection (Agilent Technologies, Santa Clara, CA, U.S.A.) was used to quantify simple sugars and organic acids. The mobile phase was water, with a flow rate of 0.5 mL/min. A Cation H Bio-Rad Micro-Guard column (304.6 mm) followed by an Aminex HPX-87H carbohydrate analysis column (300 × 7.8 mm) (Bio-Rad Laboratories, Inc.; Hercules, CA, U.S.A.) was used to separate organic acids and simple sugars isocratically at 35°C. The injection volume of the sample was 10 μL and the run time was 15 minutes.

The chromatographic data were acquired, monitored, and processed using OpenLAB CDS LC Chemstation M8307AA. All compounds were identified through co-injection of authentic standards. Quantification was based on external calibration curves constructed for seven standard solutions for sugars (glucose, fructose, arabinose, galactose, and sucrose) and organic acids (lactic-, acetic-, and citric acid) in concentrations ranging from 0.05 – 10 g/L.

### 2.8 Free amino acid analysis

Samples (section 2.6) were freeze-dried (Lyovapor L-200, Buchi) for two days. To each freeze-dried sample, 1 mL of 0.1 M HCl was added, and the mixture was ultrasonicated (Branson 2210, Emerson) for five minutes. To the 200 μL sonicated sample, 200 μL of an ice-cold solution of 12.5% trichloroacetic acid was added. After incubation for 2 hours at 4°C, the sample was centrifuged at 10,000 g for 20 minutes. The supernatant was neutralized with 45 μL of freshly prepared 1 M NaOH and then mixed with 195 μL of 50 μM aminocaproic acid (internal standard). Free amino acids were quantified using HPLC as described in Rehlund et al. (2025). All proteinogenic free amino acids were quantified, except cysteine and proline, using external calibration curves. HPLC data were pre-processed using Chromeleon 7.2.7 (Thermo Scientific, MA, USA).

### 2.9 Quantification of cereulide produced by strain Y48H10 using HPLC-MS/MS

Quantification of cereulide from emetic strain Y48H10 was performed in two biological duplicates. Four technical replicates were pooled together for each sampling time point. Cereulide analysis was performed using ISO 18465 standard (ISO18465:2017, 2017) method with few modifications. Briefly, samples were diluted in 1:10 ratio with HPLC-grade acetonitrile (VWR Chemicals, Poland). Labelled cereulide (^13^C6-cereulide) was added as an internal standard in each sample at the concentration of 10 ng/ml. The cereulide was extracted by a rotating shaker (300 rpm) for 1 hour followed by resting for 30 minutes at ambient temperature. The solution was decanted twice (first 4500 *g* for 15 minutes and second 12,000 *g* for 15 minutes). A 200 µL of the supernatant was transferred into HPLC vial for LC- MS/MS analysis.

Samples were randomized and analysed on an Elute UPLC system coupled to an Impact II QTOF-MS instrument (Bruker Daltonik, Germany). Chromatographic separation was performed on a reverse-phase Supelco Discovery C18 column (10 cm × 2.1 mm, 5 µm particle size) maintained at 36 °C. The mobile phase consisted of an isocratic elution with 5% phase A (10 mM ammonium formate in water containing 0.1% formic acid) and 95% phase B (acetonitrile with 0.1% formic acid). The flow rate was set to 0.4 mL/min with an injection volume of 5 µL. Mass spectra were acquired in the range of 1000–1300 *m/z* at 12 Hz using electrospray ionization (ESI) in positive-ion mode. Prior to analysis, the instrument was externally calibrated with 10 mM sodium formate in 50% isopropanol and tuned using the ESI-L Low Concentration Tuning Mix (Agilent Technologies, CA, USA). ESI parameters were as follows: end plate offset, –500 V; capillary voltage, 4500 V; nebulizer pressure, 2.2 bar; dry gas flow, 10 L/min; and drying temperature, 220 °C. MS settings included funnel 1 and 2 RF, 200 Vpp; hexapole, 50 Vpp; ion energy, 5 eV; collision energy, 5 eV; and pre-pulse storage, 5 µs. Peak areas of natural cereulide and isotope-labelled cereulide were extracted using corresponding precursor ions at m/z 1170–1171 and 1176–1177, respectively.

### 2.10 Data analysis

Isothermal microcalorimetric data were analysed from 1 to 20 hours (BHI medium) or 24 hours (food) after inoculation, as the first hour was needed for thermal equilibration of the instrument. Baseline correction was applied using the heat flow data collected before plate insertion. For 2 out of 96 experiments, an alternative baseline heat flow value was selected to achieve a better fit after plate insertion at 50 minutes. Data from one replicate experiment corresponding to strain *B. subtilis* PRO31 was excluded, due to negative heat flow values at the beginning of the experiment, which was attributed by the instrument manufacturer to moisture evaporation from the vial.

The heat curves (heat accumulated as a function of time) were fitted with the modified Gompertz model to estimate microbial growth characteristics (Zwietering et al., 1990), based on two assumptions (1) that cells remain in the medium, and (2) that heat flow data present themselves with one central peak or a closely similar shape (e.g. peak with a shoulder) (Braissant et al., 2010). Data analysis of the isothermal calorimetry was conducted using RStudio (Posit team, 2025). Curve fitting and parameter estimation were performed on each experiment using the grofit R package, based on Gompertz modelling, to estimate the maximum heat production (mJ), the length of the calorimetric lag time (h), and the maximum calorimetric growth rate (mJ · h^−^¹) (Kahm et al., 2010). The lag time was corrected for the one-hour pre-incubation period.

Statistical tests were performed using the Gompertz parameters related to species, with a significance level of 0.05. Group comparisons between *B. cereus* and *B. subtilis* and between the interspecies clades in BHI medium were performed using unpaired T-tests, and phylogenetic comparisons between *B. cereus* clades were performed using Kruskal-Wallis with Dunn’s post-hoc test. Comparison between heat curves obtained in BHI media and food matrices of selected *B. cereus* (Y48H10, SDA GR177, MC67) and *B. subtilis* strains (PRO99, DSM10, DSM21393) were conducted using paired t-test with Bonferroni correction.

Gompertz parameters were square-root transformed to ensure equal variance among groups. A two-way analysis of variance (ANOVA) was performed to examine the effects of *Bacillus* clade (*B. cereus* and *B. subtilis*), food (milk, oat, pea, and oat-pea), and their interaction on the square-root transformed Gompertz parameters. A separate two-way ANOVA was performed to test the influence of all matrices (foods and BHI medium), *Bacillus* clade, and their interaction on square-root transformed Gompertz parameters. All ANOVA models were fitted using the lmerTest package, which extends lme4 by providing p-values for mixed effects. Post-hoc analyses were conducted using estimated marginal means (EMM) via the emmeans package.

For the metabolomics data, a three-way ANOVA was performed to evaluate the influence of *Bacillus* clade, metabolic time-point (peak vs. end), and food matrix, as well as their interactions, on the content of individual sugars, individual organic acids, and total free amino acids. Biological replicates were included as a random effect using lme4, and significance testing was carried out with lmerTest. Post-hoc analyses were conducted using EMM. Additionally, a two-way ANOVA was conducted to investigate the effects of fermentation (non-fermented vs. fermented) and matrix on total sugars. Tukey post-hoc comparisons were performed between non-fermented and fermented samples within each matrix (oat, pea, and oat–pea). Non-detected compounds were assigned a value equal to half the lowest detected concentration to enable statistical comparisons. Principal component analysis (PCA) was applied to investigate potential interactions between the metabolites of *Bacillus* spp. in food matrices using RStudio (R Core Team, 2023), and a biplot was created using the ggplot2 package.

The quantification of cereulide produced by *B. cereus* strain Y48H10 in food matrices was based on mass spectrometric peak area ratios derived from baseline-corrected chromatograms, using a signal-to-noise ratio (SNR) threshold of 3. The limit of detection (LOD) and limit of quantification (LOQ) were determined from the standard deviation of the noise in 0.1 ng/mL samples, which represented the lowest concentration at which cereulide was not detectable. Inspection and processing of obtained cereulide data were performed using MATLAB 2023b.

## 3. Results

### 3.1 Taxonomy and phylogeny

A maximum-likelihood phylogenetic tree was constructed using the UBCG pipeline (Na et al., 2018) from the whole-genome sequences of the *B. cereus* and *B. subtilis* strains (Figure 2). Given the extensive recent updates to *B. cereus* taxonomy (Carroll et al., 2022), species names in the phylogeny reflect the closest taxonomic matches based on ANIm analyses. These best-match designations are used throughout the Results section, whereas Table 1 lists them together with the earlier, previously accepted taxonomy. For strains BC59 and LWL3, previously classified as *B. cereus*, ANIm values exceeded the species threshold (>96%) for both *Bacillus wiedmannii* and *Bacillus pretiosus*. However, we adopted *B. wiedmannii* as the taxonomic designation, in line with (Mozhaitseva et al., 2025), who reported genomic similarity between these species and grouped them within a shared cluster. Moreover, *B. pretiosus* remains an effectively published but not validly recognized name under current bacterial nomenclature rules (Oren et al., 2023; Robas Mora et al., 2023), further supporting the use of *B. wiedmannii*.

**Figure 2:**
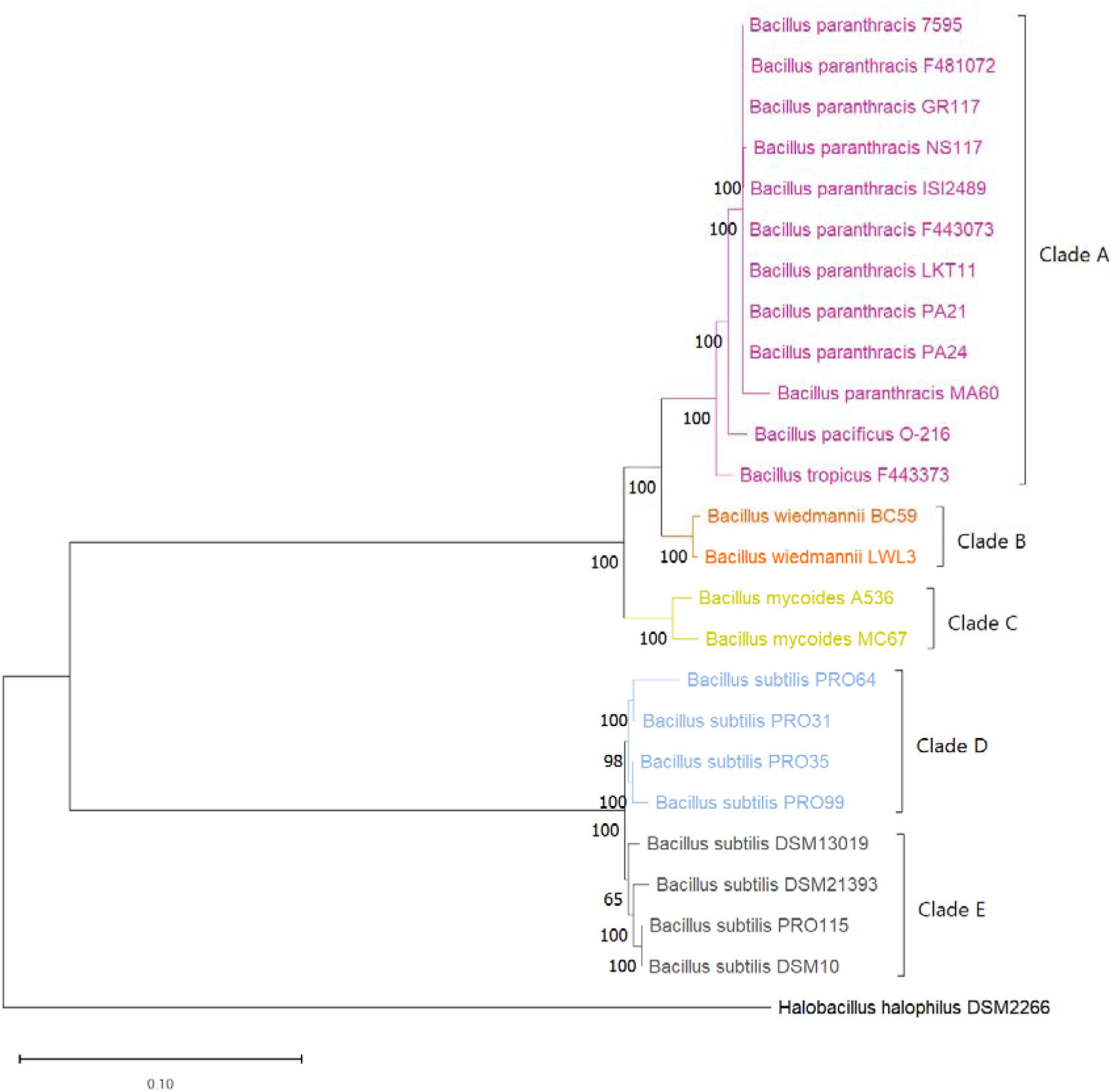
Whole genome phylogeny shows that the *Bacillus cereus* group strains cluster into three clades (A–C), while *B. subtilis* strains form two distinct clades (D–E). Rooted maximum likelihood phylogenetic tree of *Bacillus cereus* group strains (Clades A, B, and C) and *Bacillus subtilis* strains (Clades D and E) using the up-to-date bacterial core gene (UBCG) pipeline, drawn by MEGA11. The names of the species were drawn from the best taxonomic (ANIm) matches, as displayed in Table 1. *Halobacillus halophilus* DSM2266 was used to root the tree. Percentage bootstrap values are indicated at branching points, and the scale bar represents 0.01 substitutions per site.

Five different clades were identified (Figure 2), differentiating the *B. cereus* group strains into Clade A (*B. paranthracis, B. pacificus,* and *B. tropicus*), Clade B (*B. wiedmannii/B. pretiosus*), and Clade C (*B. mycoides*), and distinguishing the *B. subtilis* group into Clade D (PRO31, PRO35, PRO64, PRO99), and Clade E (DSM13019, DSM21393, PRO115, and DSM10). In short, we observed a diverse range of *B. cereus* (encompassing five different genomic subspecies) and *B. subtilis* isolates (two distinct clades) to facilitate comparisons between phylogeny and microcalorimetric analysis of metabolic activity.

### 3.2 Metabolic activity of *Bacillus* cultivated in BHI and food matrices

To investigate the metabolic activity of *Bacillus* strains under controlled conditions, we monitored the metabolic heat production profiles of 16 *B. cereus* strains and 8 *B. subtilis* strains. Cultivation was first carried out in BHI medium at 30°C for 20 hours, until the metabolic activity reached a steady-state plateau (Figure 3B). During 20 hours of cultivation, *B. cereus* and *B. subtilis* exhibited a typical sigmoidal pattern in the heat production curves (Figure 3A). The heat flow profiles of both *B. subtilis* and *B. cereus* exhibit one main peak followed by a second shoulder peak (Figure 3B).

**Figure 3:**
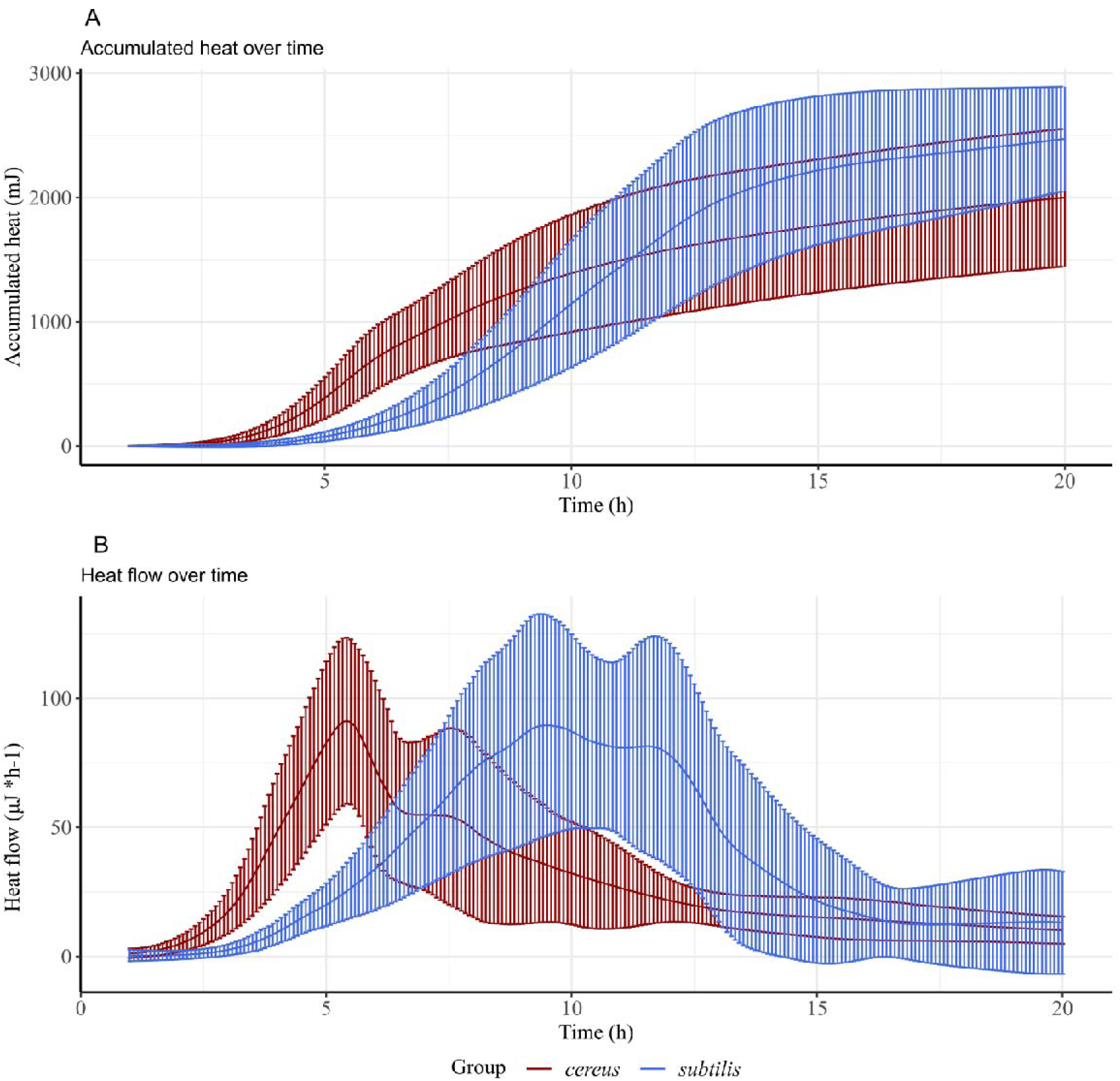
Accumulated-heat and heat-flow data indicate a delayed onset of detectable metabolic activity in *B. subtilis* compared with *B. cereus*, with *B. subtilis* ultimately generating more total heat. Metabolic heat characteristics of 16 *B. cereus* (red) and 8 *B. subtilis* (blue) strains cultivated in BHI medium, based on four biological replicates across two different experiments (A) Mean values (curve) and standard deviation (error bars) of accumulated heat production over time (B) Mean values (curve) and standard deviation (error bars) of heat flow over time

To gain a better understanding of the metabolic heat profiles, fitted parameters from the Gompertz model, calorimetric lag time (h), the maximum calorimetric growth rate (mJ*h^−1^), and total heat (mJ), were plotted for each *Bacillus* group (*cereus* vs. *subtilis*) (Figure 4A) and the phylogenetic clades within *B. cereus* and *B. subtilis* groups (A, B vs. C; D vs. E) (Figure 4B). Between the *B. subtilis* and *B. cereus* groups, significant differences were observed across all parameters. *B. subtilis* released a significantly higher amount of total heat (2732 mJ ± 462) than *B. cereus* (1941 mJ ± 566) (p < 10^−10^). *B. subtilis* displayed a higher maximum metabolic activity (378 mJ*h^−1^ ± 140) than *B. cereus* (240 mJ*h^−1^ ± 106) (p < 10^−5^). Finally, a longer lag time was observed for *B. subtilis* (7.9 hours ± 1.0) than for *B. cereus* (4.3 hours ± 0.8) (p < 10^−16^).

**Figure 4:**
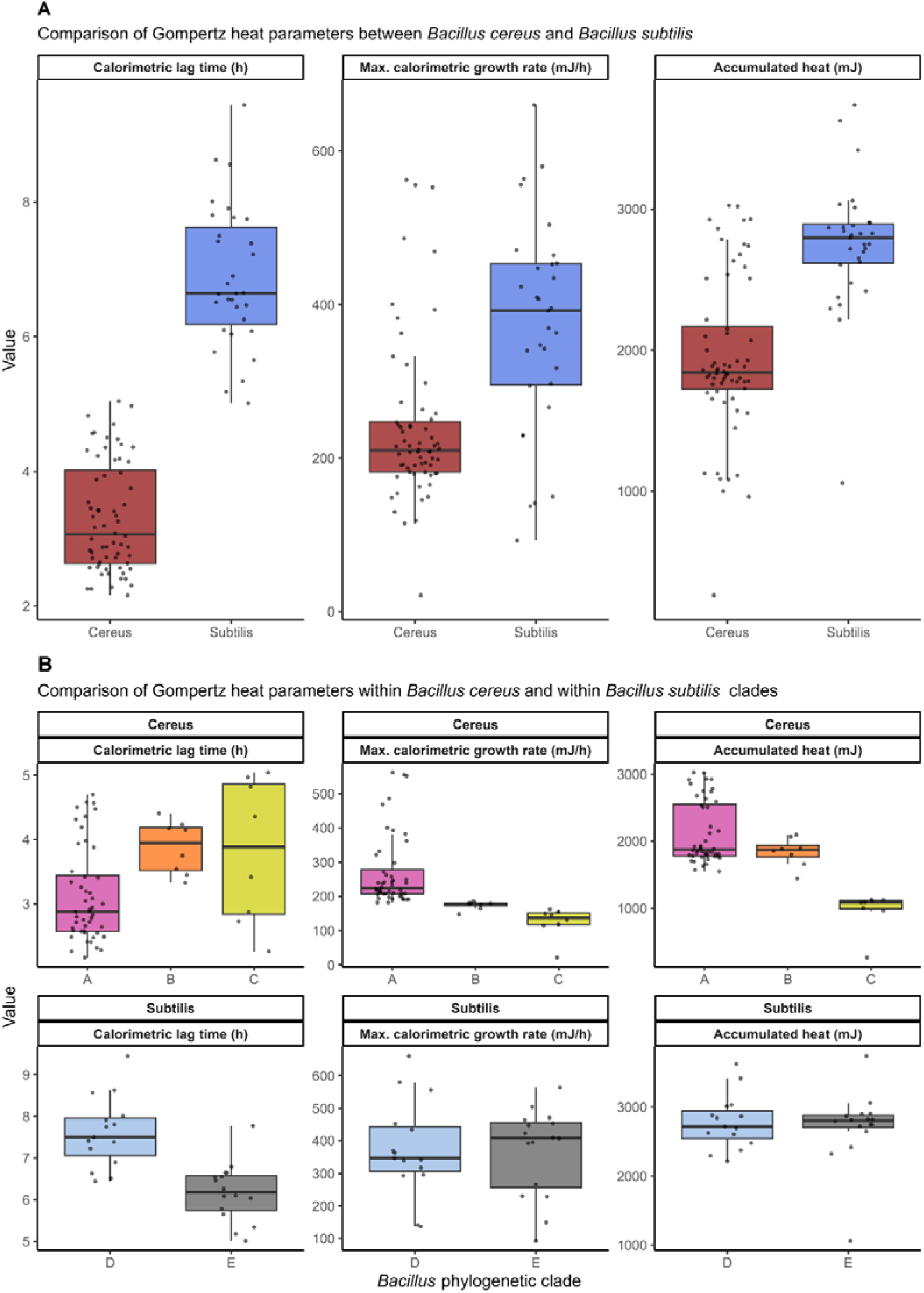
All Gompertz parameters derived from the heat-flow curves differed significantly between *B. cereus* and *B. subtilis*, with additional parameter-specific variation observed within clades. Calorimetric lag time (h), maximum calorimetric growth rate (mJ*h^−1^), and total heat (mJ) after 20 hours. Calorimetric lag time was corrected for 1 h pre-incubation period. Each dot represents an individual biological replicate (four replicates per *Bacillus* strain) within each phylogenetic group. (A): Differences between the *B. subtilis* and *B. cereus* groups. (B): Differences among the phylogenetic clades as defined in Figure 2, between Clade A (*B. paranthracis, B. pacificus*, and *B. tropicus*), B (*B. wiedmannii*), C (*B. mycoides*), D (*B. subtilis* PRO31, PRO35, PRO64, PRO99), and E (*B. subtilis* DSM13019, DSM21393, PRO115, and DSM10)

In addition, differences in metabolic activity were observed within the *B. subtilis* and *B. cereus* groups. Between the *B. cereus* clades, *B. mycoides* strains from clade C produced a significantly lower amount of total heat (971 mJ ± 292) than *B. paranthracis*, *B. pacificus,* and *B. tropicus* strains from clade A (2119 mJ ± 464, p <10^−5^) and *B. wiedmannii* strains from clade B (1839 mJ ± 212, p = 0.01). In addition, the maximum calorimetric growth rate differed significantly between Clade A and B (p <10^−5^) and between Clade A and C (p <10-5). Between the *B. subtilis* clades, clade D strains exhibited a longer lag time (8.6 hours ± 0.8) than Clade E (7.2 hours ± 0.7) (p <10^−5^).

In addition to BHI, three strains of *B. cereus* (Y48H10, SDA GR177, MC67) and three *B. subtilis* (PRO99, DSM10, DSM21393) were cultured in milk (pH 6.5), oat (pH 7.1), pea (pH 6.9), and oat-pea (pH 5.8) at 30°C. Their metabolic heat activity was monitored for 24-hour periods. Gompertz parameters were subsequently derived from the calorimetric data (Table 2).

**Table 2:** Gompertz calorimetric growth parameters (mean (± sd)) of *B. cereus* (Y48H10, SDA GR177 and MC67) and *B. subtilis* (PRO99, DSM10 and DSM21393) cultivation in milk, oat, pea, and oat-pea matrices.

| Species | Food | Calorimetric lag time [hours] | Max calorimetric growth rate [$\text{mJ}\cdot\text{h}^{-1}$] | Max accumulated heat [mJ] |
| --- | --- | --- | --- | --- |
| *B. cereus* | Milk | 7.4 (± 0.5) | 150 (± 8) | 2040 (± 120) |
| | Oat | 14.5 (± 12.8) | 184 (± 97) | 1696 (± 865) |
| | Pea | 7.7 (± 0.3) | 216 (± 65) | 1663 (± 714) |
| | Oat-pea | 9.8 (± 1.7) | 407 (± 103) | 1880 (± 665) |
| *B. subtilis* | Milk | 12.4 (± 2.7) | 109 (± 30) | 2057 (± 334) |
| | Oat | 11.6 (± 3.1) | 211 (± 43) | 2452 (± 212) |
| | Pea | 10.4 (± 1.3) | 275 (± 40) | 2065 (± 301) |
| | Oat-pea | 10.9 (± 0.4) | 419 (± 28) | 2403 (± 232) |

*B. subtilis* exhibited a more consistent calorimetric lag time across food matrices, with a longer lag time in milk (12.4 hours ± 2.7) compared to *B. cereus* (7.35 hours ± 0.5). Maximum calorimetric growth rates varied significantly by substrate and were higher in the combined oat-pea matrix compared to oat (p = 0.002, EMM), pea (p = 0.01, EMM), and milk (p < 10^−4^, EMM). Across all food matrices, *B. subtilis* produced more metabolic heat than *B. cereus* (p = 0.02, EMM) after 24 hours, consistent with observations on BHI medium after 20 hours. When comparing *Bacillus* metabolism between BHI and the food matrices, the calorimetric lag time was significantly lower in BHI than in oat (p = 0.02, EMM), while no significant differences were observed between the other matrices.

### 3.3 Metabolic profiling of *Bacillus* growth in oat, pea, and oat-pea matrices

Targeted metabolomic profiling was performed to assess how matrix composition influences *Bacillus* metabolic activity, with a particular focus on the enhanced metabolic peak observed in the combined oat-pea matrix. One *B. subtilis* (DSM21393) and one emetic *B. cereus* strain (Y48H10) were selected for cultivation on the food matrices and sampled at two time-points: at peak metabolic activity, e.g. at maximum heat flow (8 hours for *B. cereus,* 11 hours for *B. subtilis*) and at minimum metabolic activity (24 hours for both).

Sugars, organic acids, and free amino acids were quantified in all samples. To interpret the multifactorial data generated from targeted metabolomic profiling (simple sugars, organic acids, and free amino acids), principal component analysis (PCA) was employed to visualize trends in metabolite synthesis and consumption by *B. subtilis* and *B. cereus* cultured in plant-based matrices over 24 hours. PCA revealed that matrix composition was the main driver of sample separation (Figure 5). PC1 (41% variance) was associated with free amino acids, galactose, and fructose. In comparison, PC2 (25% variance) separated the pea matrix from the oat and oat-pea, reflecting lower sugar levels (sucrose, glucose) and higher citrate content.

**Figure 5:**
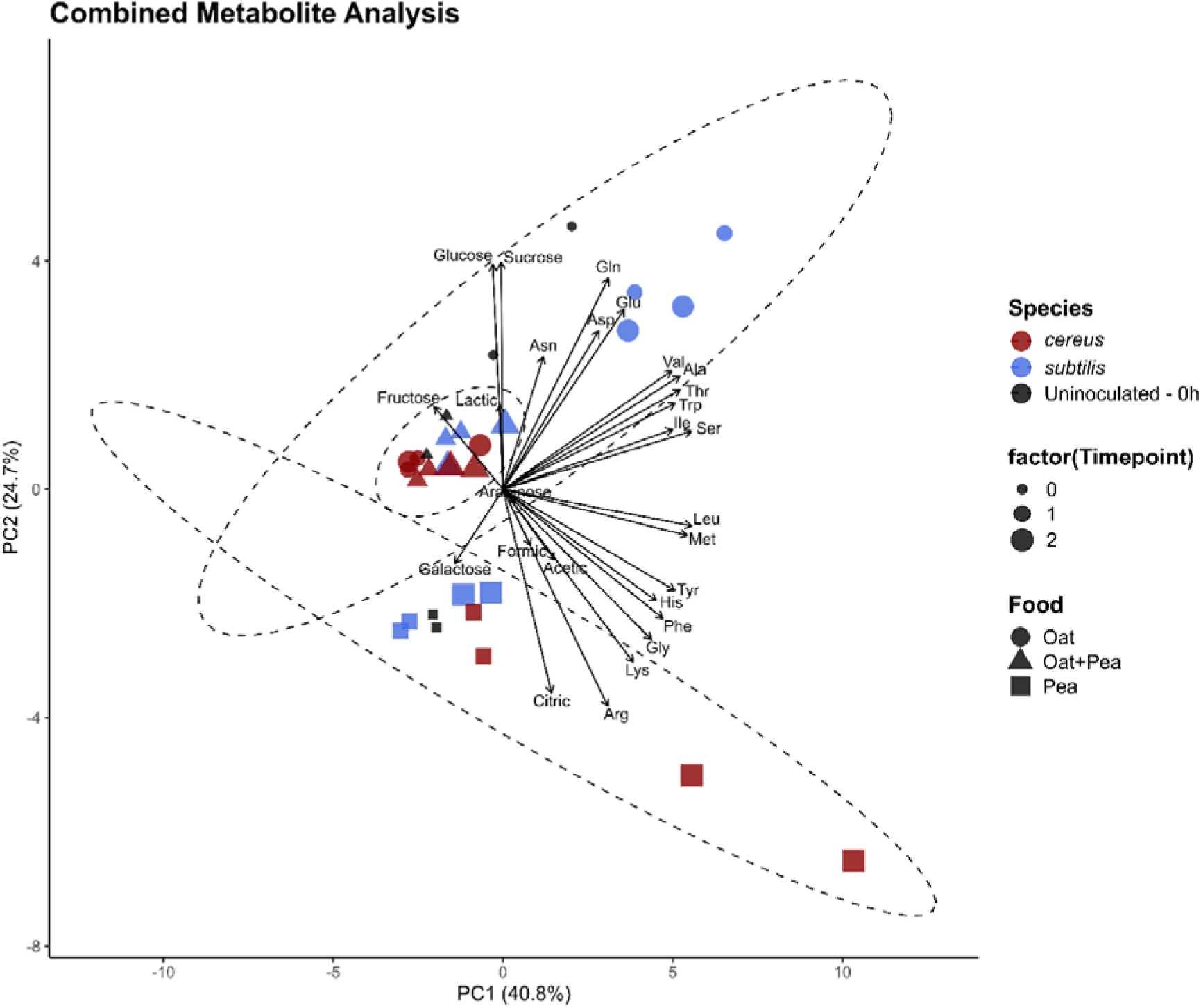
PCA of sugars, organic acids, and free amino acids shows separation of pea matrices from oat and oat-pea, and illustrates metabolic shifts associated with *Bacillus* cultivation. PCA biplot of metabolites from *Bacillus cereus* Y48H10 (red) and *Bacillus subtilis* DSM21393 (blue) cultivated in oat (circles), pea (squares), and oat-pea (triangles) with two biological replicates. Metabolomic profiles of non-inoculated food matrices are visualized in black. Ellipses represent 95% of the population for each food matrix (pea, oat, oat-pea).

To gain insight into the factors driving PCA separation, we analysed temporal changes in sugars, organic acids, and free amino acids. Matrix composition strongly influenced sugar and organic acid profiles. Total sugar content, as quantified by HPLC-RID, was significantly higher in the uninoculated oat (49.4 ± 1.8 g/L) and oat-pea (52.6 ± 3.6 g/L, p = 0.001, Tukey, Figure 6A) matrices compared to pea (0.74 ± 0.00 g/L, p = 0.001, Tukey, Figure 6A). *Bacillus* fermentation significantly reduced the total amount of sugars in the oat-pea matrix (p = 0.01, two-way ANOVA, EMM).

**Figure 6:**
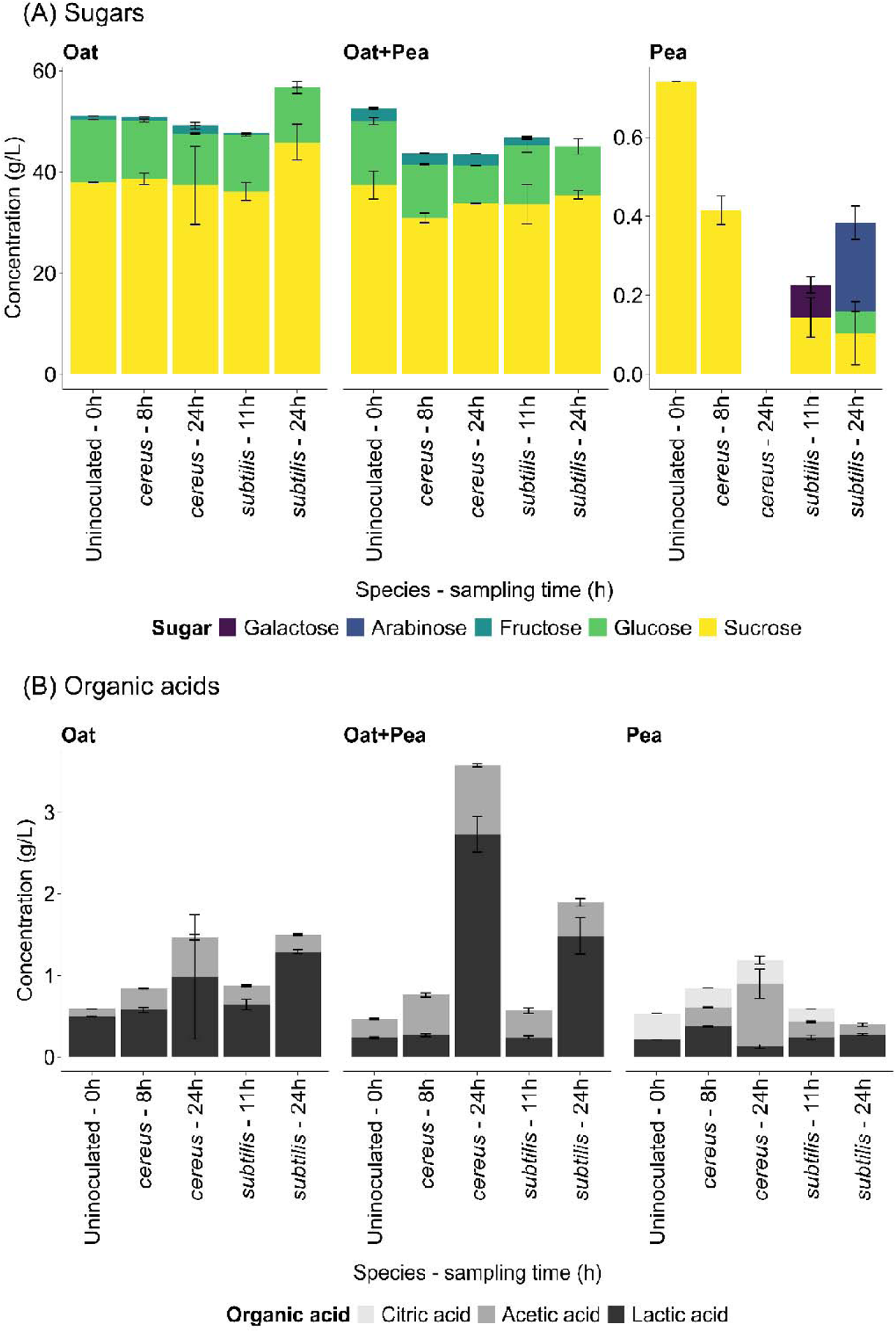
Sugar and organic acid profiles shows matrix-dependent and *Bacillus*-mediated changes, including significant sugar depletion and organic acid accumulation in the oat–pea matrix. Metabolic profiling of *Bacillus cereus* Y48H10 and *Bacillus subtilis* DSM21393 cultivated in oat, pea, and oat-pea matrices. Error bars are based on two biological replicates, each of which consists of four pooled technical replicates. The stacked bars represent the total sugars (A) and organic acids (B), with each segment showing the contribution of individual compounds.

Citrate was uniquely present in the pea matrix and absent from both oat and oat-pea (Figure 6B), contributing to its distinct separation in PCA (Figure 5). Cultivation of *B. subtilis* and *B. cereus* led to an increase in total organic acids from peak metabolism (maximum heat flow) to end metabolic phase (p = 10^−5^, three-way ANOVA). The oat-pea matrix showed the most pronounced increase in organic acids, with both species sharply increasing lactic acid levels (p = 10^−4^, EMM, Figure 6B). In contrast, the changes in organic acid levels in oat and pea were more modest (p > 0.05, EMM). During *B. subtilis* growth on pea, citrate was consumed, and acetic acid was produced. *B. cereus* produced more acetic acid than *B. subtilis* after 24 hours (p = 10^−4^, EMM).

Free amino acids were observed in all uninoculated matrices, with the highest levels detected in the (non-autoclaved) oat drink (Figure 7). *B. cereus* and *B. subtilis* had distinct effects on the free amino acid profiles depending on the matrix. The most pronounced effects were observed during *B. subtilis* growth on oat and *B. cereus* growth on pea, both increasing free amino acid levels, as also reflected in the PCA (Figure 5). These effects differed in the types of free amino acids released and the timing of their accumulation. For *B. cereus* in pea, total free amino acids increased from 59 µmol/g ± 1 (8 hours) to 219 µmol/g ± 54 (24 hours) (p = 0.003, EMM), primarily due to accumulation of Gly, Phe, Ala, Ser, Leu and Tyr. In contrast, *B. subtilis* displayed higher total free amino acid levels in oat than in pea (p = 10^−4^, EMM), peaking at 227 µmol/g ± 38 at maximum metabolic activity after 11 hours, with elevated levels of Asp, Gly and Ala, and a reduction in Asn. In the oat-pea matrix, *Bacillus* did not significantly alter total free amino acid levels over time (p = 1, EMM).

**Figure 7:**
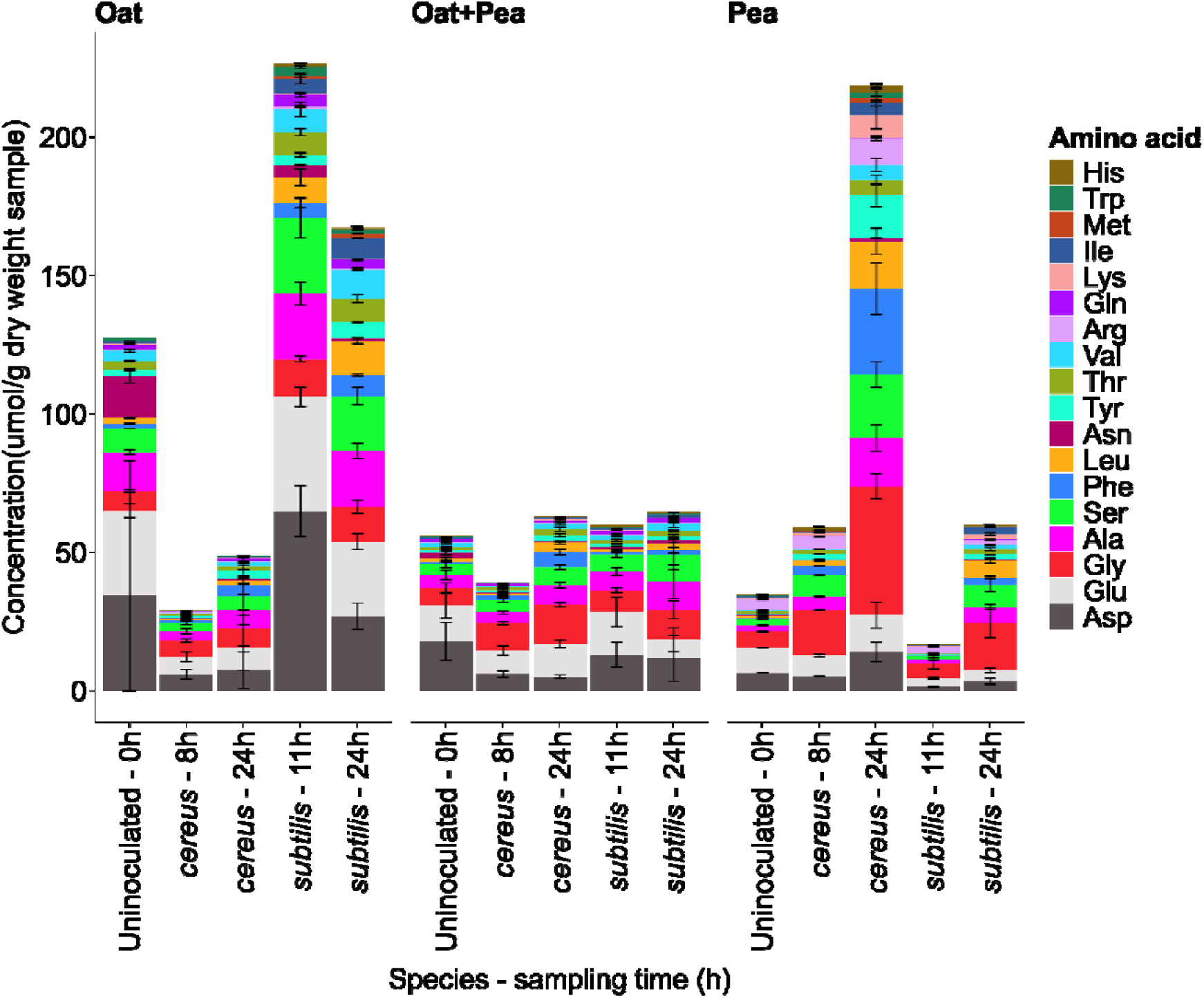
Free amino acid profiles showed species- and matrix-specific responses: *B. cereus* strongly increased total amino acids in pea, *B. subtilis* generated the highest amino acid levels in oat, and neither species substantially changed total amino acid content in oat–pea. Free amino acid profile of *Bacillus cereus* Y48H10 and *Bacillus subtilis* DSM21393 cultivated in oat, pea, and oat-pea matrices. Error bars are based on two biological replicates, each consisting of four pooled technical replicates. The stacked bars represent the total free amino acids, with each segment showing the contribution of individual amino acids.

Cereulide was exclusively quantified in the uninoculated matrices and in the *B. cereus* samples, since it was assumed that *B. subtilis* is incapable of producing cereulide. Cereulide was below the detection limit (0.1 ng/mL) in the pea matrix. In contrast, concentrations of 3.0 and 2.9 ng/mL were detected in the oat-pea matrix, while the oat drink exhibited substantially higher concentrations of 46 and 48 ng/mL in the two biological replicates.

## 4. Discussion

To examine the metabolic heat production of *B. cereus* and *B. subtilis*, we analyzed 24 *Bacillus* strains representing diverse origins, pathogenic potential, and genetic backgrounds. The pathogenic potential - reflected in the ability to produce enterotoxins and/or emetic toxins (Table 1) - partially aligned with phylogenetic clustering, suggesting a link between genetic lineage and toxin capacity. All emetic strains showed the closest ANIm resemblance to *B. paranthracis*, except the emetic psychrotolerant strain *B. mycoides* M67. Enterotoxin-producing strains were identified in all three *B. cereus* clades and were not distinguishable by phylogeny or IMC-derived metabolic profiles. Strain classification within the *B. cereus* group (Table 1) reflects recent taxonomic revisions. *B. wiedmannii* was proposed as a new species in 2016 (Miller et al., 2016). *B. paranthracis, B. pacificus, and B. tropicus* were proposed in 2017 (Liu et al., 2017), and *B. weihenstephanensis* was merged with *B. mycoides* in 2018 (Liu et al., 2018). Although a formal species-level taxonomic framework is still needed for the *B. cereus* group, whole-genome phylogeny supports inference of clade-dependent metabolic behaviour.

Clade-specific growth dynamics were evident across all Gompertz parameters during cultivation in BHI at 30°C. *B. subtilis* exhibited a higher model-estimated total heat output (2745 mJ) than the observed value (2473 mJ), indicating sustained metabolic activity beyond the 20-hour endpoint. In contrast, *B. cereus* reached an earlier plateau, with modelled (1941 mJ) and measured (2000 mJ) values closely aligned. Both species displayed a shoulder peak following the main heat flow peak (Figure 3B), likely reflecting a shift in substrate utilization. Calorimetric lag phase differed markedly: *B. cereus* averaged 4.3 h ± 0.8, significantly shorter than *B. subtilis* (7.9 h ± 1.0), with no overlap.

These metabolic growth profiles likely reflect distinct ecological strategies in nutrient-rich media such as BHI. *B. subtilis* may sustain metabolic activity and produce more heat in BHI via a broader peptide/carbohydrate catabolism supported by a diverse enzymatic repertoire. In contrast, *B. cereus* showed rapid onset and early saturation of metabolic activity, which may reflect a faster environmental adaptation and a less energetically intensive growth strategy in nutrient-rich conditions. Phylogenetic variation within each group further influenced metabolic heat production. *B. mycoides* (clade B) strains released significantly less heat than other *B. cereus* clades. One member of clade B, MC67, is psychrotolerant (Thorsen et al., 2006), which likely exhibits optimal activity below 30°C which may explain the reduced metabolic heat output detected for clade B at this incubation temperature.

Clade-specific metabolic differences, including calorimetric lag time, may depend on initial inoculum size. Lankford et al (1966) demonstrated that increasing *Bacillus* inoculum size shortens the cell-based lag phase, presumably because a larger proportion of cells are already in a physiologically active state (Lankford et al., 1966). In isothermal microcalorimetry, detection time also depends on inoculum size, with Fricke et al. (2019) showing a log-linear relationship between initial cell concentration and time-to-detection in *Lactobacillus plantarum* (Fricke et al., 2019). Importantly, calorimetric lag time does not necessarily equal cell-based lag time, as IMC detection depends on reaching the signal threshold. Documented IMC detection limits vary widely, e.g. *B. subtilis* has been detected from as low as < 10 CFU per vial (Brueckner et al., 2016), whereas *Proteus mirabilis* required ∼10 cells per vial (Braissant et al., 2015b), while standardized assessments of IMC sensitivity across taxa are still lacking. Consequently, the IMC-based lag times we observed in our study (*B. cereus* 4.3 h ± 0.8, *B. subtilis* 7.9 h ± 1.0) reflect when metabolic activity became detectable, and should be interpreted in the context of the inoculum used (OD600 0.2, ∼10^6^ CFU/mL)(Fricke et al., 2019).

The influence of substrate composition on *Bacillus* metabolic kinetics became more apparent when strains were cultivated in food matrices. To test the persistence of clade-specific metabolic traits across matrices, three *B. cereus* strains and three *B. subtilis* strains were cultivated in milk, oat, pea, and combined oat-pea matrix. As in BHI medium, *B. subtilis* generally exhibited longer lag times than *B. cereus*, most notably in milk (12.4 hours vs. 7.4 hours). However, species-level distinctions were less pronounced in the food matrices compared to BHI, suggesting that substrate composition exerted a stronger influence on physiological responses than phylogenetic background.

To elucidate the biochemical mechanisms underlying matrix-dependent growth, we profiled sugar, organic acids, and free amino acids during cultivation of one representative *B. cereus* and *B. subtilis* strain in oat, pea, and oat-pea. Direct comparison of metabolic profiles across matrices was complicated by differential thermal processing. The oat drink had already undergone UHT processing and was not autoclaved, as autoclaving inhibited *B. cereus* growth. Autoclaving of oat drink likely degraded heat-sensitive nutrients such as free amino acids (Evans & Butts, 1949), which are known to support *B. cereus* growth (Canoy et al., 2025; Agata et al., 1999). However, autoclaving the matrices containing pea was necessary to solubilize the proteins and eliminate contaminants. This discrepancy introduces an important limitation, as thermal treatment history becomes a confounding variable. Standardized thermal processing would help ensure observations reflect matrix differences and not the treatment, as heat can trigger Maillard reactions, deplete key nutrients, and alter the redox balance (Lund & Ray, 2017). These effects likely contributed to the elevated free amino acid levels observed in the non-autoclaved oat matrix compared to the autoclaved pea-containing matrices.

Despite these limitations, clear matrix-dependent metabolic trends emerged. The oat-pea matrix supported the highest *Bacillus* activity, showing the highest maximum calorimetric growth rate for both *Bacillus* species across all tested matrices and substrates (Table 2). This suggests that combining oat and pea provides a more favourable nutrient environment than either substrate alone. Consistent with this, *Bacillus* spp. accumulated the highest levels of organic acids in the oat-pea matrix, including lactic and acetic acids (Figure 6B). These responses suggest that complementary interactions between peptides and free amino acids from pea and readily fermentable sugars from oat could promote more efficient carbohydrate metabolism, possibly by optimizing carbon–nitrogen balance and enhancing glycolytic flux, and thereby contributing to higher calorimetric growth. This interpretation aligns with findings by Chen et al. (2023), who reported that *B. subtilis* strains grew faster in medium containing meat extract supplemented with glucose than with meat extract alone, with glucose addition increasing acetate formation under well-aerated conditions and lactate accumulation under oxygen limitation, reflecting enhanced carbon flux toward organic acids. Similar nutrient-driven enhancement has been demonstrated in acidogenic fermentations involving microbial communities composed of multiple bacterial genera, where mixed ratios of soluble carbohydrates and proteins resulted in higher levels of organic acids and yields compared with single-nutrient substrates (Vazquez-Fernandez et al., 2025).

Because pH can also influence microbial metabolism, we assessed whether initial acidity might contribute to the enhanced activity observed in the oat–pea system. The autoclaved oat-pea matrix had a lower initial pH (5.8) than autoclaved pea (pH 6.9) and non-autoclaved oat matrix (pH 7.1). Autoclaving likely contributed to these pH differences, since we observed a similar pH decrease when UHT oat drink was subjected to autoclaving, from pH 7.1 to 5.3. Given comparable organic acid levels among the non-inoculated matrices (Figure 6B), the pH drop likely reflects abiotic heat-induced glucose degradation (Leitzen et al., 2021), especially in the carbohydrate-rich oat matrices. However, moderately lower pH alone is unlikely to explain enhanced metabolic activity in the oat-pea matrix. In *B. subtilis,* moderate acidic conditions (pH 6.0) upregulate acid-stress pathways rather than promoting further acidification (Wilks et al., 2009). Similarly, in *B. cereus*, low pH has been associated with reduced production of acid fermentation end products and shifts toward alternative metabolites under stress, rather than increased acidogenesis (Le Lay et al., 2015). Together, these results suggest that enhanced *Bacillus* metabolism and organic acid accumulation in the oat–pea matrix is primarily nutrient driven, although pH-nutrient interactions cannot be entirely ruled out.

To further dissect matrix compositional effects, we focused on the pea matrix, characterised by low sugar content, the unique presence of citrate, and clear PCA separation from oat and oat-pea. During cultivation on pea, both *B. cereus* and *B. subtilis* produced and consumed organic acids, but their metabolic patterns diverged. *B. cereus* produced more acetic acid than *B. subtilis* at the end of metabolism (24 hours). *B. subtilis* significantly reduced citrate, consistent with its ability to take up citrate via Mg² -citrate transporters CitM/CitH and use it as a carbon source (Schilling et al., 2007; Warner & Lolkema, 2002). In contrast, citrate levels remained unchanged during *B. cereus* growth in pea. Citrate is rarely reported as a carbon source for *B. cereus*, although specific strains can metabolize it (Venkateswaran et al., 2017). These species-specific responses suggest that sugar limitation in pea may drive differential organic acid turnover.

Species-specific strategies also emerged in free amino acid dynamics, revealing differential nutrient interactions of *B. cereus* and *B. subtilis*. *B. subtilis* showed transient free amino acid release in oat, consistent with its capacity to metabolize complex carbohydrates and synthesize amino acids under nutrient-rich conditions. In contrast, *B. cereus* accumulated free amino acids in pea, likely reflecting proteolytic activity and amino acid catabolism under nutrient or oxygen stress. This interpretation aligns with the inherently low-oxygen environment of the calorimetric ampoules, where samples are enclosed with only a small headspace, naturally restricting oxygen transfer. Limited changes of free amino acid levels in the oat-pea matrix, despite high organic acid production, suggest a shift away from amino acid turnover toward carbohydrate fermentation. These observations align with previous findings that *B. cereus* preferentially utilizes proteinaceous substrates under limited oxygen conditions (Chang et al., 2021).

Cereulide, the heat- and acid-stable emetic toxin produced by *B. cereus*, was quantified across food matrices. Cereulide was detected in oat and oat-pea matrices, but not in pea, indicating that composition influences toxin production. Notably, although *B. cereus* Y48H10 exhibited the highest heat production in the oat-pea matrix, cereulide levels were lower than in oat alone, indicating that toxin synthesis does not necessarily correlate with overall metabolic activity. This pattern aligns with previous studies showing that *B. cereus* can grow comparably well in diverse foods, including bechamel sauce, pasta, and potato puree, while generating highly variable cereulide levels (Rajkovic et al., 2006). Likewise, Ellouze et al. (2021) observed faster cereulide formation by *B. cereus* F4810/72 in a cereal-based matrix (rice) than in a vegetable matrix (vegetable purée) despite slower bacterial growth in rice, underscoring the matrix dependency of cereulide production. In contrast to their vegetable matrix, we detected no quantifiable cereulide in pea at 30°C within 24 hours, indicating that some vegetable-based matrices may not provide the nutritional or ecological conditions required for toxin biosynthesis.

Together, these observations highlight that cereulide formation is driven more by specific environmental and nutritional cues than by growth alone. Cereulide expression is highly sensitive to pH, nutrient profile, and redox state (Dommel et al., 2010; Ehling-Schulz et al., 2015). While pH is known to affect cereulide production, no clear association was observed between organic acid levels (Figure 6) and cereulide levels, suggesting that acidification alone could not explain the differences in cereulide accumulation between the matrices. Consistent with previous food-based studies (Marxen et al., 2015; Agata et al., 2002; Messelhäusser et al., 2014; Rajkovic et al., 2006), we observed the highest cereulide concentrations in the carbohydrate-rich matrix (oat). This supports the idea that carbohydrate-rich food environments may trigger a dysregulated metabolic response, leading to higher cereulide production than in the bacterium’s natural niche (Ehling-Schulz et al., 2015).

Building on this ecological and nutritional context, a key contribution of this study is the controlled comparison of cereulide production in three matrices with distinct nutrient profiles: a carbohydrate-rich oat matrix, a nitrogen-rich pea fraction, and a mixed oat-pea matrix. The absence of detected cereulide in pea and the reduced levels in oat-pea relative to oat suggest that carbon–nitrogen balance and/or amino acid availability may influence the regulatory pathways or precursor supply required for cereulide biosynthesis. This interpretation is supported by Agata et al. (1999), who found that amino acids strongly influence growth and cereulide production and that high concentrations (50mM) of leucine, isoleucine and glutamic acid reduce cereulide formation. Consistent with this, *B. cereus* grown in the pea matrix markedly increased free amino acid levels, including isoleucine and leucine (Figure 7). Collectively, these observations suggest that pea-derived constituents alter nutrient cues in ways that disrupt regulatory networks governing *ces* expression or limit precursor pools needed for non-ribosomal peptide synthesis. Whether these effects represent active suppression of biosynthesis or primarily constrained precursor availability remains unresolved and warrants further investigation.

While cereulide is most frequently associated with starchy foods (Messelhäusser et al., 2014), outbreaks also occur in legume-based products such as fermented soybeans (Inatsu et al., 2020) and soaked navy beans (Nicholls et al., 2016), indicating that low-carbohydrate environments can support toxin synthesis under permissive conditions. Thorsen et al. (2011) found no emetic strains in a survey of African bread snacks made from legumes, whereas Kyrylenko et al. (2023) reported that 9% of *B. cereus* isolates from plant-based dairy alternative ingredients carried cereulide-producing genes. In the context of this variability, our findings underscore that nutrient composition, particularly the availability of fermentable carbohydrates and their interaction with nitrogen sources, plays a central role in determining cereulide production potential across plant-based food matrices.

Metabolic heat profiling in monoculture provided complementary insights into *Bacillus* physiology, distinguishing species- and clade-level behaviour under controlled conditions. Our findings reveal that *B. subtilis* consistently produces more metabolic heat than *B. cereus*. While calorimetric profile differences at the clade-level were clearly distinguishable in BHI, these contrasts were considerably diminished in the food matrices, highlighting the limitations of directly transferring laboratory media observations to complex substrates like food. Although calorimetry alone cannot resolve metabolic pathways, its integration with metabolomic profiling offers a robust framework to assess microbial activity, adaptation, and potential safety risks in complex food systems. While the metabolic capacities of *B. subtilis* and *B. cereus* are well-characterized in defined media, their proteomic responses in food matrices remain underexplored (Yeo et al., 2012). Future studies integrating calorimetry, metabolomics, and proteomics could help reveal matrix-specific regulatory mechanisms that govern spoilage activity and beneficial fermentation traits in *B. subtilis,* as well as toxin production in *B. cereus*. Such approaches are essential to understand *Bacillus* behaviour across diverse food environments.

## 5. Conclusion

This study demonstrates that *Bacillus cereus* and *Bacillus subtilis* exhibit distinct metabolic behaviour across food-relevant matrices. In BHI at 30°C with an initial inoculum of 10^6^ CFU/mL, *B. subtilis* showed longer calorimetric lag times and higher total heat production than *B. cereus*. However, in food matrices, substrate composition had a more pronounced effect on metabolic activity than species identity, although *B. subtilis* consistently generated more heat.

The combined oat-pea matrix supported the highest calorimetric growth rates, likely driven by enhanced carbohydrate metabolism, as evidenced by reduced sugar levels and increased organic acids. In contrast, the pea matrix, characterised by low sugar levels and the unique presence of citrate, triggered post-metabolic consumption of citrate by *B. subtilis*. Free amino acid profiles reflected matrix-specific strategies, *B. subtilis* released free amino acids in oat, while *B. cereus* predominantly did so in pea. Interestingly, despite high calorimetric activity, the oat-pea matrix showed limited change in free amino acid levels, suggesting a shift toward carbohydrate-driven metabolism.

Cereulide production by *B. cereus* was strongly matrix-dependent, with high levels detected in oat, moderate levels in oat-pea, and none in pea, highlighting the role of substrate composition in cereulide synthesis. Together, these findings underscore the need for matrix-specific microbial risk assessment tools. They also demonstrate that integration of isothermal microcalorimetry with metabolomics provides a more nuanced understanding of microbial behaviour in complex food environments. Similar heat profiles can mask divergent metabolic strategies; thus, this combined approach offers a valuable tool for evaluating spoilage and contamination risks, particularly in the development of safer plant-based food products.

## Acknowledgment

The authors wish to acknowledge the Villum Foundation (HLR:34434) and the joint alliance grant between the Danish Dairy Research Foundation, Arla Foods amba, Novonesis, and the University of Copenhagen (SK: SaFeMix) for financial support. Author T.S.C. was supported by the Novo Nordisk Foundation (NNF21OC0066330).

## Author contribution

Emilie V. Skriver: Conceptualization, Methodology, Formal analysis, Investigation, Visualization, Writing – original draft, Writing – review and editing; Tessa S. Canoy: Conceptualization, Methodology, Formal analysis, Investigation, Visualization, Writing – original draft, Writing – review and editing; Yuandong Sha: Formal analysis, Investigation, Visualization; Morten A. Rasmussen: Software, Formal analysis; Bekzod Khakimov: Methodology, Formal analysis, Supervision; Susanne Knøchel: Writing – review and editing, Resources, Supervision, Project administration, Funding acquisition; Henriette L. Røder; Conceptualization, Methodology, Resources, Writing – review and editing, Supervision, Project administration, Funding acquisition.

